# Single-stride exposure to pulse torque assistance provided by a robotic exoskeleton at the hip and knee joints

**DOI:** 10.1101/582866

**Authors:** Robert L. McGrath, Fabrizio Sergi

## Abstract

Robot assisted gait retraining is an increasingly common method for supporting restoration of walking function after neurological injury. Gait speed, an indicator of walking function, is correlated with propulsive force, a measure modulated by the posture of the trailing limb at push-off. With the ultimate goal of improving efficacy of robot assisted gait retraining, we sought to directly target gait propulsion, by exposing subjects to pulses of joint torque applied at the hip and knee joints to modulate push-off posture. Our previous work investigated changes in joint moments associated with push-off posture modulation, which informed the composition of 16 joint torque pulse conditions.

In this work, we utilized a robotic exoskeleton to apply pulses of torque to the hip and knee joints, during individual strides, of 16 healthy control subjects, and quantified the effects of this intervention on hip extension and propulsive impulse during and after application of these pulses.

We observed significant effects in the outcome measures primarily at the stride of pulse application. Specifically, when pulses were applied at late stance, we observed a significant increase in propulsive impulse when knee and/or hip flexion pulses were applied and a significant increase in hip extension angle when hip extension torque pulses were applied. When pulses were applied at early stance, we observed a significant increase in propulsive impulse associated with hip extension torque.

## I. INTRODUCTION

Robot assisted gat training (RAGT) is an increasingly common tool for supporting restoration of walking ability after neurological injury [1], [2]. It offers various benefits over conventional therapy, such as reduced burden on physical therapists and the ability to objectively and accurately measure a patients performance [1], [3]. While RAGT has shown promising results [2], the efficacy of RAGT has yet to to exceed that of conventional therapy [4]. We speculate that this is due to limitations of the previously implemented robotic controllers. Some previously implemented robotic controllers include kinematic goals such as foot reference trajectory [5]–[7] which have the disadvantage of imposing relatively large constraints on the kinematics of gait. Simpler assistance strategies, such as those which utilize repetitive joint torque pulses, have been implemented with desirable effects such as entrainment of gait and reduced metabolic cost [8]–[10] while minimally constraining gait kinematics.

Moreover, previous approaches to RAGT have not directly targeted mechanisms that are associated with improvement in walking function. Gait speed (GS) is a primary outcome measure of walking rehabilitation studies, as it is associated with a better quality of life [11]. GS is known to be correlated with the propulsive force of the foot against the ground, also known as the anterior ground reaction force [12]. Propulsive impulse, the propulsive force integrated over time, is determined by two factors: ankle moment and posture of the trailing limb at push-off [13]. More recent work has determined that push-off posture has a much greater relative contribution to propulsive force than ankle moment [14]. Push-off posture can be quantified by a scalar value, trailing limb angle (TLA), defined as the angle of the line connecting the hip joint center and foot center of pressure at the instant of peak propulsive force, relative to the laboratory vertical axis [14]. However, it is currently unknown if robot-assisted modulation of the push-off posture will modulate the propulsion mechanism and associated measures of walking function. In an effort to improve efficacy of RAGT, we sought to formulate a controller designed specifically to target push-off posture and modulate gait propulsion.

We previously investigated the differences in net lower extremity joint moments associated with experimentally imposed modulation of push-off posture and GS [15]. We approximated these push-off posture dependent joint torque profile differences with pulses of torque. We compiled individual subject results via torque pulse histograms and identified clustering of joint torque pulses at specific instants of the gait cycle which would approximate the difference in joint moments at multiple push-off posture conditions. At the knee, increased push-off posture was associated with extension torque and flexion torque for early and late stance, respectively. At the hip, increased push-off posture was associated with extension torque in early swing and flexion torque in late swing. However, our study has limitations associated with the purely observational nature of our analysis, which limit the direct translation of those findings to RAGT.

In this study, we composed a set of sixteen hip and knee torque pulse conditions inspired by the torque patterns associated with the modulation of push-off posture in our previous work. We applied these sixteen pulse conditions to healthy control subjects in a single-stride intervention protocol utilizing a unilateral lower extremity robotic exoskeleton which provides actuation to the hip and knee joints. Our objective was to identify the factors of torque pulse intervention that effectively modulated kinetic and kinematic gait parameters associated with propulsion. We hypothesized that the robotic application of pulse conditions corresponding to a modulation of push-off posture observed in our previous work would modulate hip extension and propulsive impulse in the corresponding direction in the stride of pulse application, as well as in the following three strides.

## II. METHODS

### A. Subjects

Sixteen healthy adults (13 males, 3 females), naive to the purpose of the study, participated in the experiment. All subjects—age (mean ± std) 24.81 ± 2.23 yr, height 177.63 ± 5.20 cm, weight 741.56 ± 83.24 N—declared to be free of orthopedic and neurological disorder affecting normal walking function. Subjects gave informed consent according to the University of Delaware IRB protocol number 929630. Subject were required to wear their own comfortable lightweight athletic clothing and shoes for the experiment.

### B. Experimental Setup

#### 1) Equipment

All data collections were performed on an instrumented split-belt treadmill (Bertec Corp., Columbus OH, USA) that measured analog force/torque data. The ALEX II robot [16], a powered unilateral lower extremity orthosis, was utilized to apply torque to the hip and knee joints of participants. The robot is suspended by rolling carriage over the instrumented treadmill, and secured in place by total locking casters, as shown in Fig. 1. For our experiment we locked both vertical rotation degrees of freedom. The ALEX II and instrumented treadmill interface with two data acquisition cards; a PCI-6221 and a PCIe-6321 (National Instruments Corp., Austin TX, U.S.). These cards are run with a custom real-time controller written in Simulink & MATLAB (MathWorks Inc., Natick MA, USA) which acquires signals from the ALEX II and instrumented treadmill and sends command signals to the ALEX II motors at 1000 Hz.

**Fig. 1:**
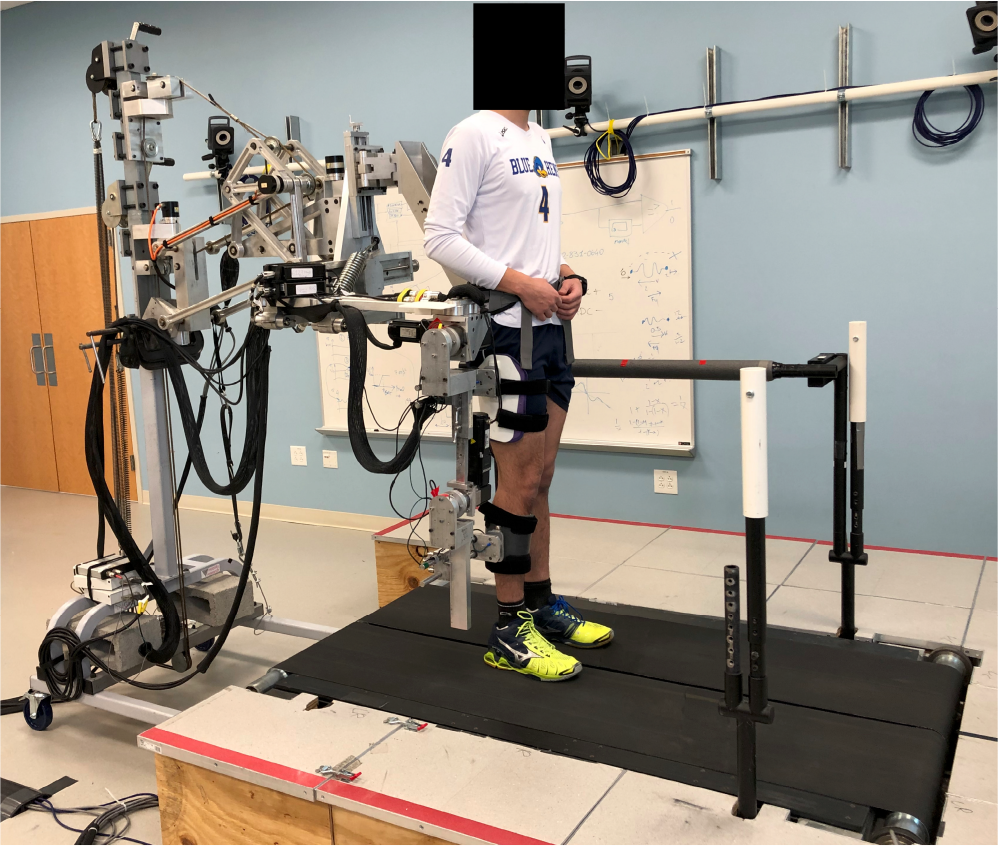
Subject wearing the ALEX II exoskeleton while walking on the instrumented treadmill.

#### 2) Controller

The ALEX II was controlled with a previously developed low-level torque-controller [17]. The controller utilized input from two 6-axis force/torque sensors located between the robot structure and the thigh/shank cuffs to compensate for the high friction in the geared motors and for the inertia of the exoskeleton structure. The highlevel controller monitors treadmill vertical ground reaction force to track right heel strikes events. These events were used to determine the start of gait cycle, and to estimate gait cycle time in real-time, based on the average of the previous six gait cycles. This estimate of gait cycle time is utilized in conjunction with pulse time values, as percentages of gait cycle, to determine timing of controller events. The selection of the torque pulse condition to be implemented by the controller determines a pulse start time, in percentage of gait cycle, amplitude in newton-meters, and a constant duration of 10% of gait cycle. The actual onset time of the torque pulse to be fed into the low-level-controller begins at 10% of gait cycle before the pulse condition specified start time in order to compensate for the delay in torque application.

### C. Experimental procedures

#### 1) Donning exoskeleton

Subjects were placed loosely in the exoskeleton and several adjustments were tuned and equipment sizes were selected prior to fastening at the waist, thigh, and shank. Final tuning was performed to ensure that each of the subjects’ feet remained on their respective treadmill belt while walking to ensure proper triggering of torque pulses to be applied to the right limb.

#### 2) Zero-torque

In zero-torque mode, the subjects walked at a slow speed which was incrementally changed until the subject specified a speed to be the fastest at which they felt comfortable. Three ramp down trials were performed, starting at their established fastest speed, until the subject reached a comfortable GS. Three ramp up trials were performed, starting at 0.6 m/s, until the subject reached a comfortable GS. These six trial GS values were averaged to determine a self selected gait speed (ss-GS) at which the subject would walk for the remainder of the experiment. Data was collected in zero-torque mode at ss-GS for a minimum of two minutes. The subject was then removed from the exoskeleton and given at least 5 minutes of rest.

#### 3) Single stride pulse sequences

Sixteen conditions of torque pulses at the knee and/or hip were tested in this experiment, as shown in Fig. 2. The pulses were square waves with a duration of 10% gait cycle and applied at a time of early or late stance, starting at 10% or 45% of gait cycle, respectively. Pulses consisted of knee extension or flexion pulses as 10 Nm or – 10 Nm in amplitude, respectively, and/or hip extension or flexion pulses as 15 Nm or – 15 Nm in amplitude, respectively.

**Fig. 2:**
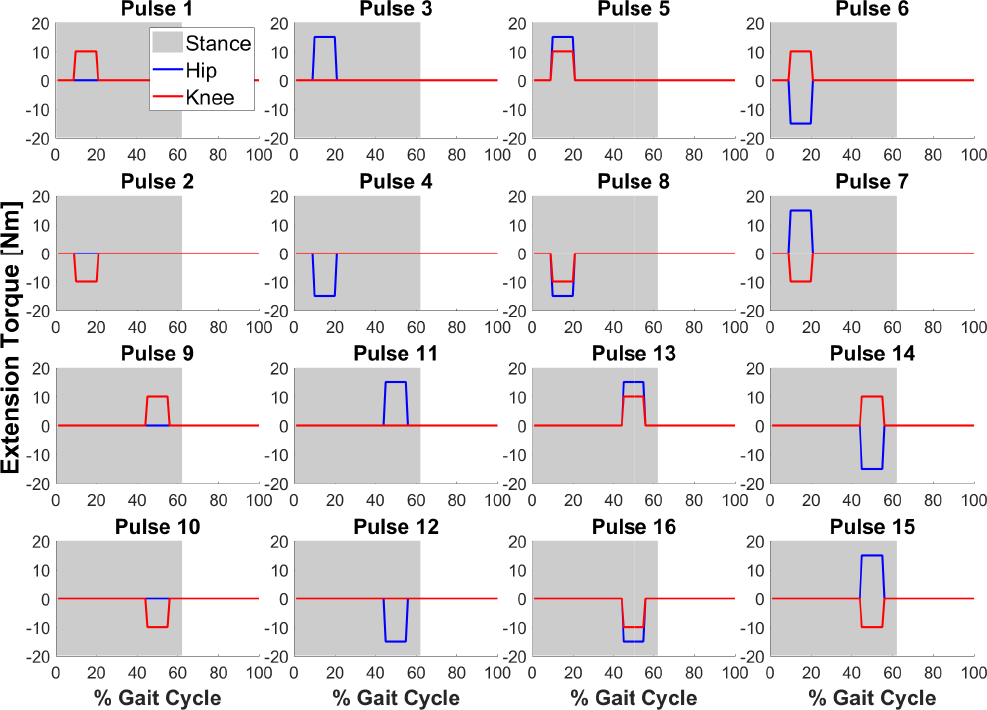
The sixteen torque pulse conditions. The two top and the two bottom rows have pulse conditions corresponding to early and late stance, respectively. First through fourth columns are knee only, hip only, knee and hip same direction, and knee and hip opposite directions, respectively.

Two separate sequences of single stride application of torque pulse conditions applied by the exoskeleton were performed while the subject walked at their ss-GS. Each sequence consisted of 5 repetitions of each of the 16 pulse conditions in a pseudo-randomized order; where each pulse was separated by 5-7 strides of no pulse application. Two 8-10 minute long repetitions of this experiment were conducted, separated by at least 5 minutes of rest.

### D. Data analysis

#### 1) Outcome measures

The first outcome measure, hip extension angle (HE), was measured at the instant of peak anterior ground reaction force. The hip motor smart feedback device transferred measures to the motor driver which utilized an emulated encoder resolution of 4096 PPR for an effective hip angle resolution of 0.00044 deg. The second outcome measure, propulsive impulse (PI), was measured as the anterior ground reaction force integrated over the time during which the ground reaction force is oriented in the anterior direction. Both measurements were taken at a total of five strides per pulse application; the stride prior to (– 1), the stride of (0), and the three strides following (1, 2, 3) pulse application. The PI and HE measure sets were sorted according to their respective pulse condition and stride and pooled across the 16 subjects and 10 repetitions. Each of the 10 corresponding repetitions were averaged together to yield a single value for statistical analyses.

### 2) Statistical testing

#### a) Pairwise comparisons

For both measures of each pulse condition, paired t-tests were performed at a group level between the baseline stride (– 1) and each of the four following strides (0, 1, 2, 3) to determine significance at a false positive rate of *α <* 0.05. Effect sizes, known as Cohen’s D, for both measures of each pulse condition were calculated as the difference between group means of the baseline stride (– 1) and pulse application stride (0) divided by the pooled standard deviation.

#### b) Mixed effect models

We sought to identify which factors of robotic torque pulses primarily influenced the selected outcome measures of PI and HE. As such, we fit both data sets to a linear mixed effects model utilizing the “fitlme” function of the MATLAB Statistics and Machine Learning Toolbox. Each data set consisted of 1280 data points (16 subjects x 16 pulse conditions x 5 strides x 1 mean value). In the linear mixed effects models, each data point was assigned a value from the fixed effects: stride number (ordinal values: – 1, 0, 1, 2, or 3), pulse time as a percentage of gait cycle (10 or 45), hip torque amplitude in newton-meters (10, 0, or – 10), and knee torque amplitude in newton-meters (15, 0, or – 15). The random effect was subject number (nominal values of 1 through 16) and was applied to the intercept of the linear mixed effect model.

A null data subset was produced for hypothetical pulses 17 and 18 with zero torque amplitudes in early and late stance, respectively. This data subset was drawn from averaging measures for pulses 1 through 8 and 9 through 16 for pulses 17 and 18, respectively, from stride – 1. This averaged data from stride – 1 was copied to the remaining four strides 0 through 3 within each of the 16 subjects. The intercept corresponded to the fixed effect values of stride – 1, pulse time 10%, knee pulse amplitude 0 and hip pulse amplitude 0. Each model therefore has 9 main effects, 28 two-way effects, 36 three-way effects, and 16 four-way effects. In the tables, pulse time, stride, hip pulse amplitude and knee pulse amplitude are represented by GC, Str, H, and K, respectively. Main effects and interaction effects significant at a false positive rate of *α <* 0.05 are reported. For purposes of examining the effects of GC, hip pulse amplitude and knee pulse amplitude on PI and HE; four-way interaction figures of pulse time x stride x knee pulse amplitude x hip pulse amplitude were examined.

## III. RESULTS

### A. Group measures

#### 1) Propulsive Impulse

Figure 3 depicts the group means for PI by stride for each of the sixteen pulse conditions. Paired comparison between the baseline stride (– 1) and the pulse application stride (0) revealed a statistically significant effect in thirteen of the sixteen pulse conditions. Early stance extension pulse conditions 1 and 3 both increased PI, while their summation in pulse 5 led to an even greater increase in PI. Early stance flexion pulse condition 2 did not lead to a change in PI while pulse condition 4 led to a small decrease in PI. The summation of these two pulses in pulse condition 8 led to a small decrease in PI. Early stance flexion and extension summation pulse conditions 6 and 7 both significantly increased PI. Late stance knee pulse conditions 9 and 10 led to a significant decrease and increase in PI, respectively, however, late stance hip pulses 11 and 12 did not lead to significant changes in PI. Late stance summation pulses 13, 14, 15, and 16 all exhibited the effects on PI according to their knee components (pulse conditions 9 and 10). As can be seen in Figure 3, only three pulse conditions had significant difference between baseline and stride 1; pulse conditions 2, 8, and 11.

**Fig. 3:**
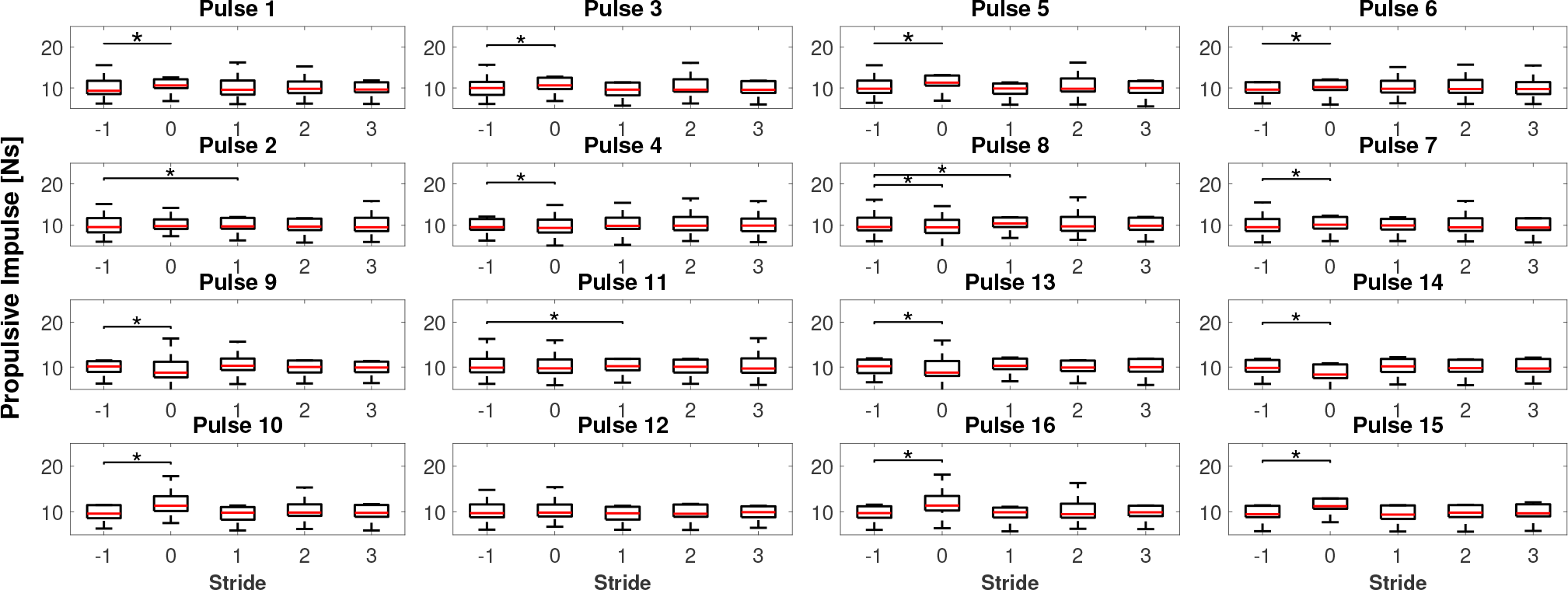
Group propulsive impulse data by stride for all pulse conditions.

#### 2) Hip Extension

Figure 4 depicts the group means for HE by stride for each of the sixteen pulse conditions and Table III summarizes the effect sizes. Paired comparison between the baseline stride (– 1) and the pulse application stride (0) revealed a statistically significant effect in nine of the sixteen pulse conditions. Early stance extension pulse conditions 1 and 3 both slightly increased HE, while their summation in pulse 5 led to an even greater increase in HE. However, early stance pulse conditions 2, 4, 6, 7, 8, did not lead to significant changes in HE. Late stance extension pulse conditions 9 and 11 increased HE and their summation in pulse condition 13 led to a large increase in HE. Late stance flexion pulse conditions 10 and 12 decreased HE and their summation in pulse conditions 16 led to a large decrease in HE. The late stance summation pulse conditions 14 and 15 did not lead to significant changes in HE. As can be seen in Figure 4, only three pulse conditions had significant differences between baseline and strides 1, 2, or 3; pulse conditions 4, 7, and 13.

**Fig. 4:**
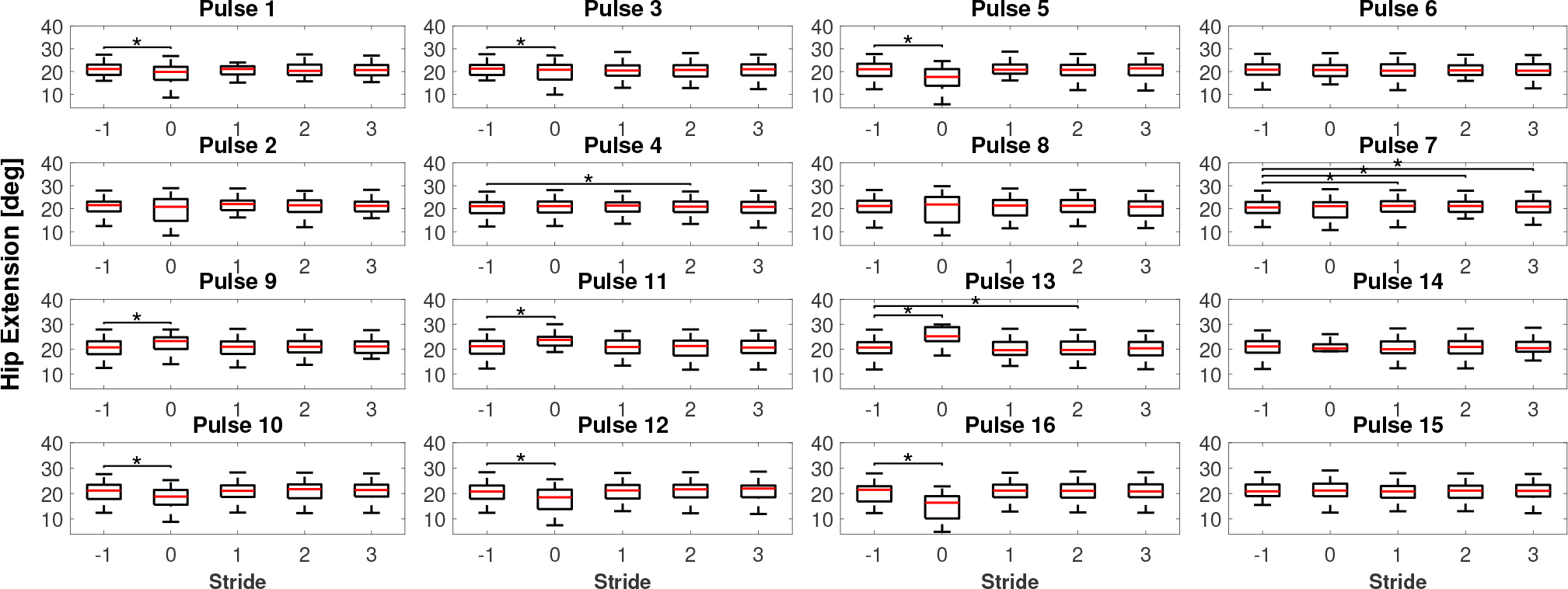
Group hip extension data by stride for all pulse conditions.

### B. Mixed effect models

#### 1) Propulsive impulse

For the PI mixed effect model, *R*^2^ ordinary equaled 0.9825 and adjusted equaled 0.9813; indicating a good fit between the data and model. The significant main effect and interaction terms are shown in Table I. As can be seen in the table, the Str_0_main effect term exists in all interactions terms, indicating that PI was effectively modulated between strides – 1 and 0 as a function of multiple experimental factors. The Str_0_⋅K_10_ interaction term indicates that knee extension torque in early stance pulse conditions increased PI. The Str_0_⋅H_*xx*_ interaction terms indicate that in early stance pulse conditions, extension torque increased PI and flexion torque decreased PI. The GC_45_⋅Str_0_⋅K_*xx*_ interaction terms indicates the late stance knee extension torque decreased PI and knee flexion torque increased PI. The Str_0_⋅K_10_⋅H_15_ interaction term establishes that knee and hip extension in early stance pulse conditions decreased PI while the GC_45_ Str_0_⋅K_10_⋅H_15_ interaction term establishes that knee and hip extension in late stance pulse conditions increased PI. As for the four-way interaction, shown in Fig. 5, depicts that in early stance pulse conditions, PI increased with hip extension torque and decreased with hip flexion torque. Instead, for late stance pulses, PI decreased with knee extension torque and increased with knee flexion torque.

**Fig. 5:**
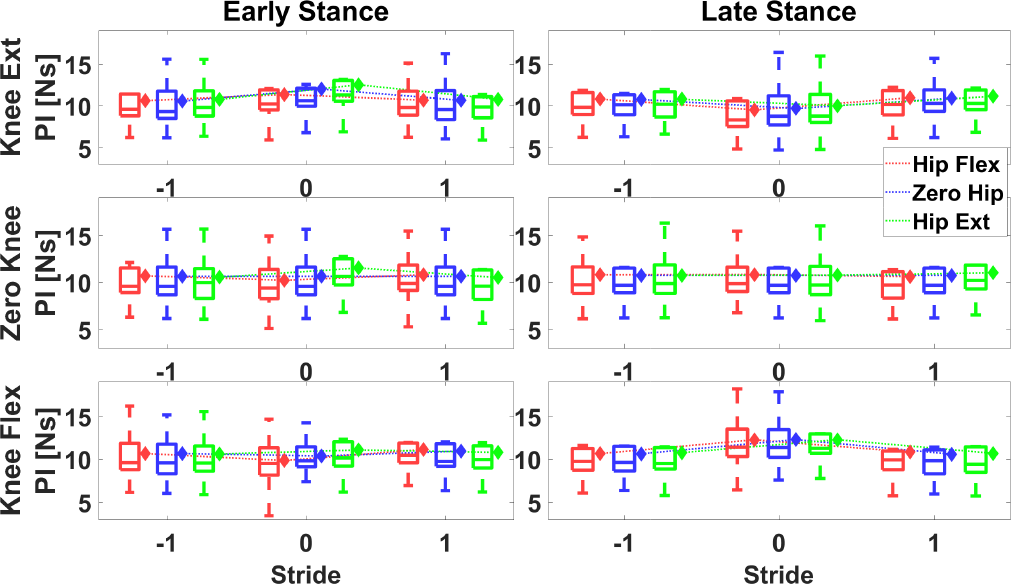
Propulsive Impulse four-way interaction. Boxplots show actual distributions of data points and diamonds are model estimated means.

**TABLE I:**
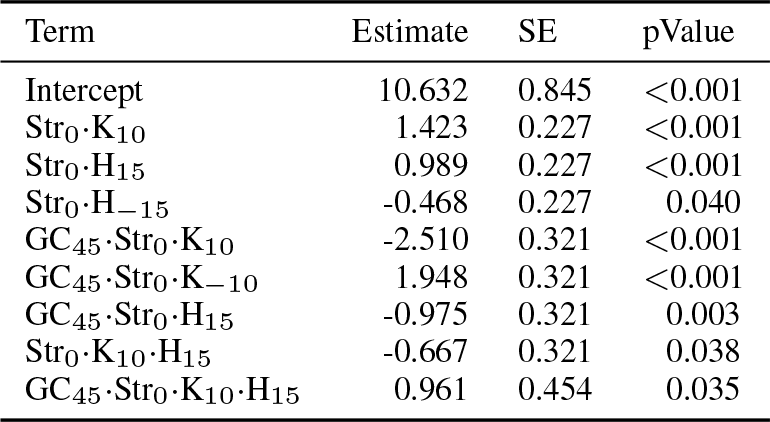
Significant results of the PI mixed effects model fitting

| Term | Estimate | SE | pValue |
| --- | --- | --- | --- |
| Intercept | 10.632 | 0.845 | <0.001 |
| Str<sub>0</sub>·K<sub>10</sub> | 1.423 | 0.227 | <0.001 |
| Str<sub>0</sub>·H<sub>15</sub> | 0.989 | 0.227 | <0.001 |
| Str<sub>0</sub>·H<sub>-15</sub> | -0.468 | 0.227 | 0.040 |
| GC<sub>45</sub>·Str<sub>0</sub>·K<sub>10</sub> | -2.510 | 0.321 | <0.001 |
| GC<sub>45</sub>·Str<sub>0</sub>·K<sub>-10</sub> | 1.948 | 0.321 | <0.001 |
| GC<sub>45</sub>·Str<sub>0</sub>·H<sub>15</sub> | -0.975 | 0.321 | 0.003 |
| Str<sub>0</sub>·K<sub>10</sub>·H<sub>15</sub> | -0.667 | 0.321 | 0.038 |
| GC<sub>45</sub>·Str<sub>0</sub>·K<sub>10</sub>·H<sub>15</sub> | 0.961 | 0.454 | 0.035 |

#### 2) Hip extension

For the HE linear mixed effect model, *R*^2^ ordinary equaled 0.9361 and adjusted equaled 0.9319; indicating a good fit between the data and model. The significant interaction terms are shown in Table II. As can be seen in the table, the Str_0_ main effect term exists in all interactions terms, indicating that HE is effectively modulated between strides baseline – 1 and application 0 as a function of multiple experimental factors. The Str_0_⋅K_*xx*_ interaction terms indicate that both knee flexion and extension in early stance pulse conditions decreased HE. The GC_45_⋅Str_0_⋅K_10_ interaction term indicates that in late stance pulse conditions, knee extension increased HE. The GC_45_⋅Str_0_⋅H_*xx*_ interaction terms establish that in late stance pulse conditions, hip extension and flexion, increased and decreased HE, respectively. The four-way interaction is shown in Figure 6. The figure shows that for late stance pulse conditions, HE decreased with hip flexion torque and increased with hip extension torque. Also in late stance pulse conditions, HE increased with knee extension and decreased with knee flexion.

**Fig. 6:**
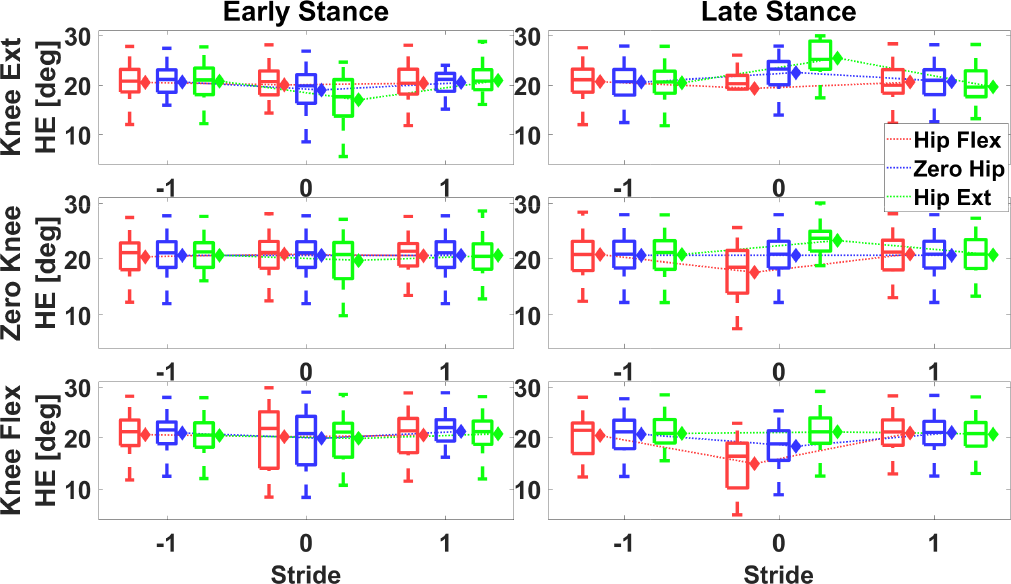
Hip Extension four-way interaction. Boxplots show actual distributions of data points and diamonds are model estimated means.

**TABLE II:**
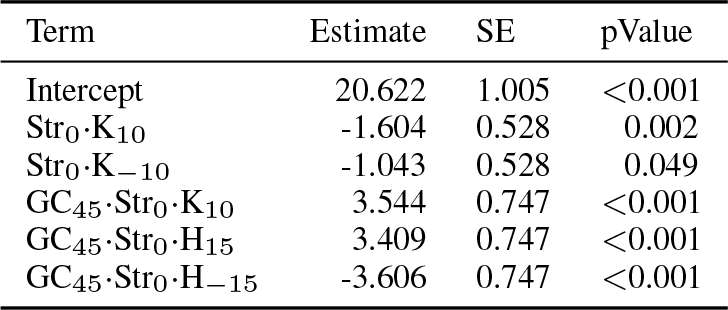
Significant results of the HE mixed effects model fitting

| Term | Estimate | SE | pValue |
| --- | --- | --- | --- |
| Intercept | 20.622 | 1.005 | <0.001 |
| Str<sub>0</sub>·K<sub>10</sub> | -1.604 | 0.528 | 0.002 |
| Str<sub>0</sub>·K<sub>-10</sub> | -1.043 | 0.528 | 0.049 |
| GC<sub>45</sub>·Str<sub>0</sub>·K<sub>10</sub> | 3.544 | 0.747 | <0.001 |
| GC<sub>45</sub>·Str<sub>0</sub>·H<sub>15</sub> | 3.409 | 0.747 | <0.001 |
| GC<sub>45</sub>·Str<sub>0</sub>·H<sub>-15</sub> | -3.606 | 0.747 | <0.001 |

## IV. DISCUSSION AND CONCLUSIONS

We formulated a set of sixteen hip and knee torque pulse conditions based on the patterns associated with push-off posture modulation observed in our previous work. We exposed healthy control subjects to these sixteen pulse conditions in a single-stride intervention protocol utilizing a robotic exoskeleton actuating about the hip and knee joints. Utilizing the sensorized robot and treadmill, we measured the modulation of HE and PI, and investigated the factors of pulse torque assistance which effectively modulated these parameters.

For group effects of the torque pulse conditions, we examined the differences between the baseline (– 1) and each of the following four strides (0, 1, 2, 3). For early stance timed pulses, conditions 1 through 8, the modulation of PI and HE were often significant but small in magnitude (Table III). The most significant modulation of these parameters were generally due to the late stance timed pulses, conditions 9 through 16. The late stance knee extension torque pulse – condition 9 – decreased PI and increased HE, while the opposing late stance knee flexion torque pulse – condition 10 – increased PI and decreased HE. The late stance hip extension torque pulse – condition 11 – did not modulate PI, but increased HE, while the opposing late stance hip flexion torque pulse – condition 12 – also did not modulate PI, but decreased HE. The late stance pulses which are summations of hip and knee torque pulses often show decoupled effects in terms of the two outcome measures. Pulse condition 14 with knee extension and hip flexion torques decreased PI but did not modulate HE. Pulse condition 15 with knee flexion and hip extension torques increased PI but did not modulate HE. Instead, pulse condition 13, with knee and hip extension torques, decreased PI and greatly increased HE. Also, pulse condition 16, with knee and hip flexion torques, increases PI and greatly decreased HE. Only a few pulse conditions yielded differences between the baseline stride (– 1) and the three strides following pulse application (1, 2, 3).

**TABLE III:** Comparison between the effect on TLA as predicted by our previous work [15], and group effect sizes for HE and PI between strides *−*1 and 0.

| Pulse | Expected effect on TLA | Effect sizes |  |
| --- | --- | --- | --- |
|  |  | HE | PI |
| 1 | Increase | -0.378 | 0.364 |
| 2 | Decrease | -0.211 | -0.079 |
| 3 | Increase | -0.208 | 0.288 |
| 4 | Decrease | 0.111 | -0.132 |
| 5 | Increase | -0.791 | 0.462 |
| 6 | Unknown | -0.121 | 0.190 |
| 7 | Unknown | -0.119 | 0.126 |
| 8 | Decrease | -0.074 | -0.221 |
| 9 | Decrease | 0.524 | -0.320 |
| 10 | Increase | -0.550 | 0.483 |
| 11 | Increase | 0.658 | 0.004 |
| 12 | Decrease | -0.678 | 0.010 |
| 13 | Unknown | 1.301 | -0.219 |
| 14 | Decrease | -0.309 | -0.370 |
| 15 | Increase | 0.071 | 0.444 |
| 16 | Unknown | -1.123 | 0.480 |

The mixed model analysis performed a break down of the effects of pulse parameters on PI and HE. Specifically, the mixed model analysis elucidated the effects of hip and knee torque during early and late stance on stride 0 as no effects existed for later strides. In early stance, PI increased with knee extension torque. Also, PI increased with hip extension and decreased with hip flexion. However, the greatest effects were measured in late stance timed pulses. Specifically, hip extension torque increased and hip flexion torque decreased HE. Also during late stance, PI decreased with knee extension torque and increased with knee flexion torque.

We had initially hypothesized that the robotic application of pulse conditions corresponding to the push-off posture modulating patterns observed in our previous work would modulate PI and HE in the same direction. In summary, the expected TLA modulations aligned well with the resulting modulations in PI, as seen in Table III. However, it is apparent in Table III that the modulation direction of HE is not consistently aligned with the direction of PI and TLA, which could be due to two factors. First, that we are only applying torque pulses for a single stride. It is possible that a single stride of intervention is not sufficient to fully elucidate the mechanistic relationship between the measures of PI, HE, and TLA. As such, repetitive stride intervention is a logical next step in further investigate the link between measures of the gait propulsion mechanism. Second, individual subject analysis not shown in this manuscript revealed high between-subject variability of the response to torque pulses. As such, to optimally modulate individual subjects’ gait, further refinement of pulse parameters is necessary. Further steps could involve human-in-the-loop optimization of pulse parameters based on various potential cost functions on an individual subject basis.

